# The genomic footprint of climate adaptation in *Chironomus riparius*

**DOI:** 10.1101/118190

**Authors:** Ann-Marie Waldvogel, Andreas Wieser, Tilman Schell, Simit Patel, Hanno Schmidt, Thomas Hankeln, Barbara Feldmeyer, Markus Pfenninger

## Abstract

The gradual heterogeneity of climatic factors pose varying selection pressures across geographic distances that leave signatures of clinal variation in the genome. Separating signatures of clinal adaptation from signatures of other evolutionary forces, such as demographic processes, genetic drift, and adaptation to non-clinal conditions of the immediate local environment is a major challenge. Here, we examine climate adaptation in five natural populations of the harlequin fly *Chironomus riparius* sampled along a climatic gradient across Europe. Our study integrates experimental data, individual genome resequencing, Pool-Seq data, and population genetic modelling. Common-garden experiments revealed a positive correlation of population growth rates corresponding to the population origin along the climate gradient, suggesting thermal adaptation on the phenotypic level. Based on a population genomic analysis, we derived empirical estimates of historical demography and migration. We used an F_ST_ outlier approach to infer positive selection across the climate gradient, in combination with an environmental association analysis. In total we identified 162 candidate genes as genomic basis of climate adaptation. Enriched functions among these candidate genes involved the apoptotic process and molecular response to heat, as well as functions identified in other studies of climate adaptation in other insects. Our results show that local climate conditions impose strong selection pressures and lead to genomic adaptation despite strong gene flow. Moreover, these results imply that selection to different climatic conditions seems to converge on a functional level, at least between different insect species.

## Introduction

Among all environmental factors, ambient temperature is most important for ectothermic organisms because it determines the rate of metabolic processes from development to reproduction (Clarke & Fraser 2004). Populations should therefore adapt to local climate and in particular to prevailing local temperatures (Clarke 2003). Moreover, recent evidence suggests temperature to also affect the speed of evolutionary processes driving population divergence (Oppold *et al*. 2016). Compared to specific environmental conditions that are restricted to a respective habitat, as e.g. anthropogenic pollutants, climate factors usually vary gradually across the globe. Accordingly, the selective pressure resulting from climate variation changes continuously, as can be expected for the evolutionary response of populations along these gradients. Through comparison of multiple populations, patterns of local adaptation should therefore be unique to single populations, whereas adaptation to climate factors should follow clinal patterns. Thus, clinal adaptation can be regarded as a special case of local adaptation that is only revealed across environmental gradients. Nevertheless, pure local adaptation can still lead to fixation of alternative alleles only at the opposite ends of a continuous environmental gradient (Barton 1999). In this case, gene flow will produce clinal patterns of allele frequency changes that do not directly result from gradual varying selection pressures. The comparison of populations from a spatially incoherent environmental gradient can thus help to avoid this genetic situation.

Evidence for clinal adaptation of phenotypic traits comes from a wide range of taxa, including plants (Loya-Rebollar *et al*. 2013; Silva *et al*. 2014), fish (local adaptation in salmonids reviewed in Fraser *et al*. 2011), invertebrates (response to thermal stress in *Daphnia*, Yampolsky *et al*. 2014), and insects in particular (Fabian *et al*. 2012; Hoffmann & Weeks 2007; Sezgin *et al*. 2004). Intensive research on *Drosophila melanogaster* populations from well-investigated environmental clines gave insight in the large- and fine-scale genomic patterns of clinal variation and its functional contribution to the adaptive phenotype (reviewed in Adrion *et al*. 2015). With single markers at first, and whole-genome data later, it was possible to identify candidate genes (e.g. the gene couch potato; Schmidt *et al*. 2008) and inversions as a form of structural genome rearrangement (Kapun *et al*. 2016) that show clinal variation in allele frequencies. However, little is known about the genomic basis of climate adaptation, particularly in non-model organisms, even though a deeper understanding is expected to bear a central role in predicting responses to global climate change (Savolainen *et al*. 2013).

The process of adaptive evolution along clines is not isolated from neutral evolutionary forces or adaptive evolution to local environmental conditions. The confounding influence of demography, introgression, and migration presents major challenges to investigate the genomic footprint of clinal adaptation (Flatt 2016). For example, Bergland *et al*. (2014; 2016) showed that clinal genetic variation could in parts be explained by an admixture of different *D. melanogaster* lineages along a latitudinal gradient. This introgression might actually be advantageous due to possible pre-adaptations of the respective ancestral lineages to their climatic conditions, highlighting the collinearity of demography and adaptation (Bergland *et al*. 2016). Moreover, endogenous genetic barriers due to disadvantageous genomic combinations that are independent of the environment can coincide with environmental boundaries and mimic the signature of local adaptation (Bierne *et al*. 2011). This illustrates the necessity to account for the potential collinearity of demography, adaptation, and endogenous genetic barriers when studying the genomic basis of clinal adaptation.

The non-biting midge *Chironomus riparius* is distributed across most parts of Europe (Pinder 1986). *C. riparius* larvae occur in small streams and ditches, predominantly in agricultural areas, and depending on the local temperature regime, the species produces multiple generations throughout the year (i.e. multivoltinism, Armitage *et al*. 1997; Loya-Rebollar *et al*. 2013; Oppold *et al*. 2016). Common-garden experiments with natural *C. riparius* populations revealed a relative fitness advantage at experimental temperatures corresponding to the respective temperature regime of their origin (Nemec *et al*. 2013). Investigating a climatic gradient from Germany to Northern Italy and Southern Spain, i.e. spatially incoherent gradient across mountain ranges, this provides a promising system for the analysis of climate adaptation in natural *C. riparius* populations.

Here, we examine temperature adaptation on the phenotypic level and integrate these results with population genomic analyses to investigate the genomic basis of climate adaptation (Figure 1) in five natural *C. riparius* populations sampled along a climatic gradient across Europe (Supporting Fig. S1.1). We conducted common-garden experiments to determine phenotypic differences in several life history traits among populations and confirmed their adaptation to different temperatures. Making use of the recently published *C. riparius* draft genome (Oppold *et al*. 2017), we performed genome-scans in a comparative outlier approach to identify signatures of adaptation. Outlier thresholds were obtained from a statistical and a modelling approach based on the demographic population histories inferred by Multiple Sequentially Markovian Coalescence (MSMC, Schiffels & Durbin 2014), and separately estimated migration rates. A spatially explicit environmental association analysis (EAA) was employed with the aim of separating the identified signatures of local from clinal adaptation.

**Figure 1:**
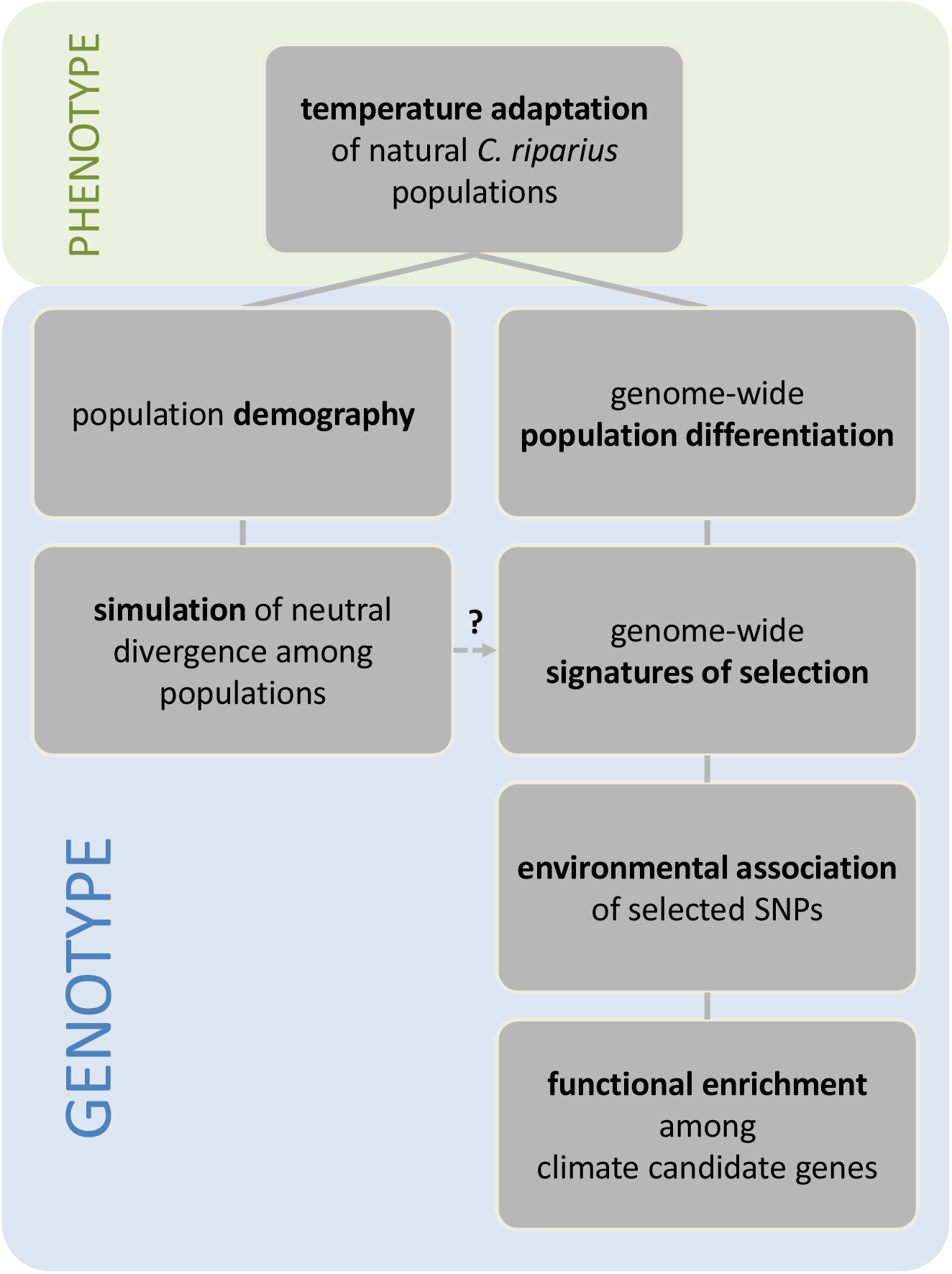
Overview of integrative approach to investigate climate adaptation on different levels of biological organisation in natural *C. riparius* populations, sampled along a climate gradient across Europe. Boxes illustrate purpose of each analysis; flow chart shows composition of results.

## Material & Methods

### Life-Cycle experiments

Phenotypic responses of natural *C. riparius* populations to different temperatures were investigated in life-cycle experiments. Populations were sampled from five locations in the field (Hesse in Germany, Lorraine and Rhône-Alpes in France, Piemont in Italy, Andalusia in Spain, see Supporting Figure S1.1 and Table S1.1) along a spatially incoherent climate gradient across the Alps and Pyrenees, and established as laboratory cultures (see Oppold *et al*. 2016 for more details). Within the first three generations in the laboratory, we tested for thermal adaptation in full life-cycle experiments at 14, 20, and 26 °C as described in Oppold et al. (2016). Phenotypic temperature adaptation was assessed by comparing population growth rates (PGR) as integrative fitness measure among populations (Supporting information 3.1). Phenotypic adaptation to different precipitation regimes is far more difficult to test experimentally and we therefore used thermal adaptation as a proxy for phenotypic climate adaptation.

### Pool-Seq data

We used 100 bp paired-end, Illumina sequencing as Pool-Seq data, derived from 105 to 168 individuals for each population to analyse allele frequency differences and gene flow among populations (see Oppold *et al*. 2017 for more details about library preparation, sequencing process, and quality processing of raw data, ENA project number PRJEB19848). Pool-Seq data was mapped to the *C. riparius* draft genome (European Nucleotide Archive accession number PRJEB15223, Oppold *et al*. 2017) and processed for downstream analysis with PoPoolation2 (v1201, Kofler *et al*. 2011, Supporting information 1.4).

### Inference of demographic population history with individual resequencing data

Demographic population history can be inferred from individual genome data using Multiple Sequential Markovian Coalescence (MSMC2, Schiffels & Durbin 2014). We therefore deep-sequenced the genomes of four randomly chosen female individuals from each population (Supporting information 4.1). Automated trimming and quality checking was performed with a wrapper script to trim the sequences (*autotrim.pl* available at https://github.com/schellt/autotrim): first round of trimming, inspection of FASTQC results, and repeated rounds of trimming if necessary to remove all overrepresented kmers, presenting partially undocumented adapter sequences. Individual resequencing data was mapped with *bwa mem* (*-M-R $individual-read-group*) and processed following the GATK best-practices pipeline (McKenna *et al*. 2010). The detailed pipeline with applied modifications is described in Oppold & Pfenninger (2017).

MSMC2 uses phased haplotype data of single chromosomes as input. We therefore prepared and phased 30 scaffolds with a minimum length of 100 kb for the analyses. Based on the annotation of the draft genome (Oppold *et al*. 2017) in combination with knowledge about the chromosomal location of certain elements or genes, we selected at least one scaffold of each of the four chromosomes. Those scaffolds covered 17.34 % of the draft genome, thus providing a representative genomic overview.

We used SHAPEIT (v2.r837, Delaneau *et al*. 2013) to phase each scaffold separately (Supporting information 4.1). Due to the absence of a *Chironomus* reference panel, all samples (i.e. four samples for each of the five populations) were merged (*bcftools merge*, v1.3, available at https://github.com/samtools/BCFtools) into one VCF containing all two-allelic SNPs for phasing. SHAPEIT accounted for the information of all 20 unrelated samples and after phasing, the output was reconverted to VCF and samples were separated again (*bcftools view-s sample-ID*).

Following the instructions for MSMC2 we used the bamCaller.py (MSMC-tools package) and the respective coverage information for each scaffold of an individual data set to generate single-sample VCF-files and mask files (Supporting information 4.1).

Mappability masks for each scaffold were generated following the procedure of Heng Li’s SNPable program (http://lh3lh3.users.sourceforge.net/snpable.shtml, accessed on 26/01/2017). MSMC2 runs, calculation of effective population size (N_e_), and cross-coalescence rates between population pairs followed the general guide of MSMC2 (available on https://github.com/stschiff/msmc/blob/master/guide.md, accessed on 26/01/2017).

### Gene flow estimation

To obtain haplotype information for multiple loci from PoolSeq data, we used the individual read information of the data (Pfenninger *et al*. 2015) and extracted 30 loci of 150 bp length, containing at least five SNPs, from the PoPoolation2 F_ST_ output file. For all loci, we considered 15 reads, thus representing haplotypic information of 15 chromosomes. We used Migrate-n (v3.6.5, Beerli 2006; Beerli & Felsenstein 2001) for Bayesian inference of the population mutation parameters (θ) and gene flow rates (number of migrants - Nm) between populations assuming a stepping-stone migration model between the nearest neighbours (MG ↔ NMF, NMF ↔ MF, MF ↔ SI, MF ↔ SS). Parameters were set as described by Pfenninger *et al*. (2015). Based on estimated θ, we calculated the expected Ne for this analysis to convert N_m_ to an effective migration rate.

### Simulation of drift expectation

In order to estimate genetic drift under neutral selective conditions and a reasonable population structure, we performed coalescent simulations using fastsimcoal2 (v. 2.5.2, Excoffier & Foll 2011). For each model we sampled 20 sequences of 1 kb length per population, applying a mutation rate per nucleotide and generation μ = 4.2 × 10^-9^ (Oppold & Pfenninger 2017), assuming no recombination and a transition bias of 0.595 (Oppold & Pfenninger 2017). The simulations ran for 200,000 iterations per model thus approximating the size of the genome. To account for different length of generation times in different populations (Oppold *et al*. 2016), we corrected N_e_ according to the respective number of generations per year (G_a_):

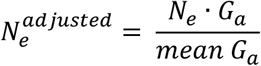

Three different demographic models were tested (Supporting information chapter 4). The basic model (*constant*) is based on a branching event of an ancestral population into five subpopulations of constant size 125,000 generations ago. Migration between neighbouring populations was allowed to varying degrees, with high migration rates from Southern France to Northern France and Southern Spain, while in accordance to the gene flow estimation, back migration is occurring more rarely (Supporting information 5).

The *growth* model retained most parameters of the previous model, adding a population growth rate of 1.0 × 10^−5^. In this model, every subpopulation accordingly expanded in size since they split 125,000 generations ago.

The final model *approximated* a demographic history with population sizes and migration rates according to the results of the MSMC analysis (see below). Population history was divided in six epochs, with symmetric migration between neighbouring populations.

Finally, we calculated pairwise F_ST_ values between each pair of simulated populations using Arlequin (v3.5, Excoffier & Lischer 2010) and compared the density functions of modelled and empirical F_ST_ values.

### Population differentiation in 1 kb-windows

Population differentiation was inferred from Pool-Seq data. SNPs from the subsampled *sync*-file were called in a sliding window approach with size of 1 kb to calculate pairwise F_ST_ values (PoPoolation2: *fst-sliding.pl* --min-count 4 --min-coverage 10 --max-coverage 51,38,67,50,45 --pool-size 336:224:210:310:236 --window-size 1000 --step-size 1000 --min-covered-fraction 0.5). We defined the upper 1 % tail of the F_ST_ distribution as statistical threshold for non-neutral differentiation (see below). All 1 kb-windows with F_ST_ above this threshold, as well as above the alternative thresholds from our simulation study (see above), were extracted for downstream analyses.

Separately, Fisher’s p-values were calculated for each 1 kb-window (PoPoolation2: *fisher-test.pl* --min-count 4 --min-coverage 10 --max-coverage 51,38,67,50,45 --window-size 1000 --step-size 1000 --min-covered-fraction 0.5). To account for multiple testing, we applied the Benjamini-Hochberg correction (Benjamini & Hochberg 1995) to all p-values (R-Core 2015). Outlier windows that remained significant after FDR correction (q < 0.01) were finally considered as highly divergent.

To investigate the genomic landscape of divergence, adjacent divergent 1 kb-windows were joined to larger divergence regions, as described in Pfenninger et al. (2015). Starting from the first outlier window encountered along a scaffold, adjacent windows were joined to an outlier region in both directions until the mean F_ST_ of the next window dropped below the cut-off value (5 % tail of the F_ST_ distribution). Then the next, not yet joined outlier window was searched and the process repeated.

### Environmental association analysis

We tested the Pool-Seq data on correlation of environmental variables with genomic differentiation using LFMM (Latent Factor Mixed Model in the frame of the ‘LEA’ R-package, Frichot & Francois 2015; Frichot *et al*. 2013) to partition clinal from local adaptation. This environmental association analysis (EAA) tool has been shown to provide the best compromise between power and error rates across various scenarios (de Villemereuil *et al*. 2014). As environmental input data, we used the first three components of a principal component analysis (PCA) with the complete set of climate data of current conditions for each location from WorldClim (Hijmans *et al*. 2005, Supporting information 2). The first three components explained 89 % of total variability and could be related to the following climatic factors (Supporting Figure S2.1-2, Table S2.1): PCA1 – gradient of cold temperatures (58 %); PCA2 – precipitation gradient (21 %); PCA3 – gradient of warm temperatures (10 %).

LFMM allows allele-frequencies from Pool-Seq data as genetic input for the EAA. To avoid spurious correlations, we included only a predefined subset of genomic loci into the analysis by considering all SNPs (SNP call with *snp-diff.pl* from PoPoolation2 with the above settings) that fell in the upper 1 % tail of the population pairwise F_ST_ distribution (estimated as described above with --window-size 1 and --step-size 1) and calculated the allele frequency of the major allele in each population for all SNPs (i.e. this includes all SNPs that exceeded the F_ST_ threshold in at least one population comparison). LFMM cannot take the pool size of Pool-Seq data into account, which reduces the power of the analysis. Referring to the Bayenv approach (Günther & Coop 2013), we artificially increased our sample size by multiplying each pooled set of frequencies with *n_i_*, where *n_i_* is the sample-size of pool *I* at this locus (i.e. in our case 20X, see above). The Pool-Seq approach assumes that each read covering a certain position originates from a different haploid chromosome. Therefore, the coverage in our data represents the number of independent chromosomes per population that were used to calculate the respective allele frequency (Futschik & Schlötterer 2010). Correspondingly, we resampled each allele frequency per population for *n_i_* = 20 from a simulated beta distribution at each locus: *beta*(*F_il_ × n_i_ + 1*, (1 − *F_il_*) × *n_i_ + 1*), with *F_il_* being the empirical frequency at locus *l* in pool *i*. This resulted in a genetic data set of 20 simulated individual allele frequencies per population at each locus. To adjust the environmental input to this resampled genetic input without creating artificial environmental variability, we only replicated each environmental factor 20 times for each locus.

We ran LFMM with five repetitions, separately for each environmental factor and a latent factor of K = 5 (i.e. number of populations, details of association tests in Supporting information Box 2.1).

Downstream analysis of the LFMM output followed the authors’ recommendations for the tool (Frichot & Francois 2015).

### Functional enrichment analysis

To define putative functional properties of differentiated genomic regions, genomic coordinates of outlier windows from population comparisons as well as clinal candidate loci identified with LFMM were compared with coordinates of protein coding genes in the *C. riparius* draft genome (Oppold *et al*. 2017). We used a custom perl-script to assign windows or loci located within a 500bp up- and downstream region of a gene. To test for gene ontology (GO) terms significantly enriched in genes lying in these regions, we intersected the lists of gene hits to produce different comparisons: gene hits of all population comparisons with gene hits from all environmental correlations (pop-pairs ~ env-variables), gene hits clinal to environmental variables amongst each other (precipitation ~ temp_cold ~ temp_warm), gene hits from outlier windows specific for one particular population for comparison against the other populations. GO terms and KEGG pathways were annotated to all protein sequences using InterProScan (v5.20-59, Jones *et al*. 2014), resulting in a complete set of terms as reference for the functional enrichment analysis. The enrichment analysis was carried out with the topGO R package (Alexa & Rahnenführer 2016) in the category ‘biological processes’, using the weight01 algorithm and Fisher statistics. Enriched GO terms were retained with a p-value of ≤ 0.05.

## Results

### Life-cycle experiments

Full life-cycle common-garden experiments revealed different responses of our natural *C. riparius* populations to the three test temperatures. Corresponding to their position on the European temperature gradient, the two Northern populations (Hesse MG and Lorraine NMF) had higher population growth rate (PGR) at 14 °C than the three remaining populations from warmer locations further South (Rhône-Alpes MF, Piemont SI, Andalusia SS), whereas this pattern was inverted at 26 °C (Fig. 2, Supporting Fig. S3.1).

**Figure 2:**
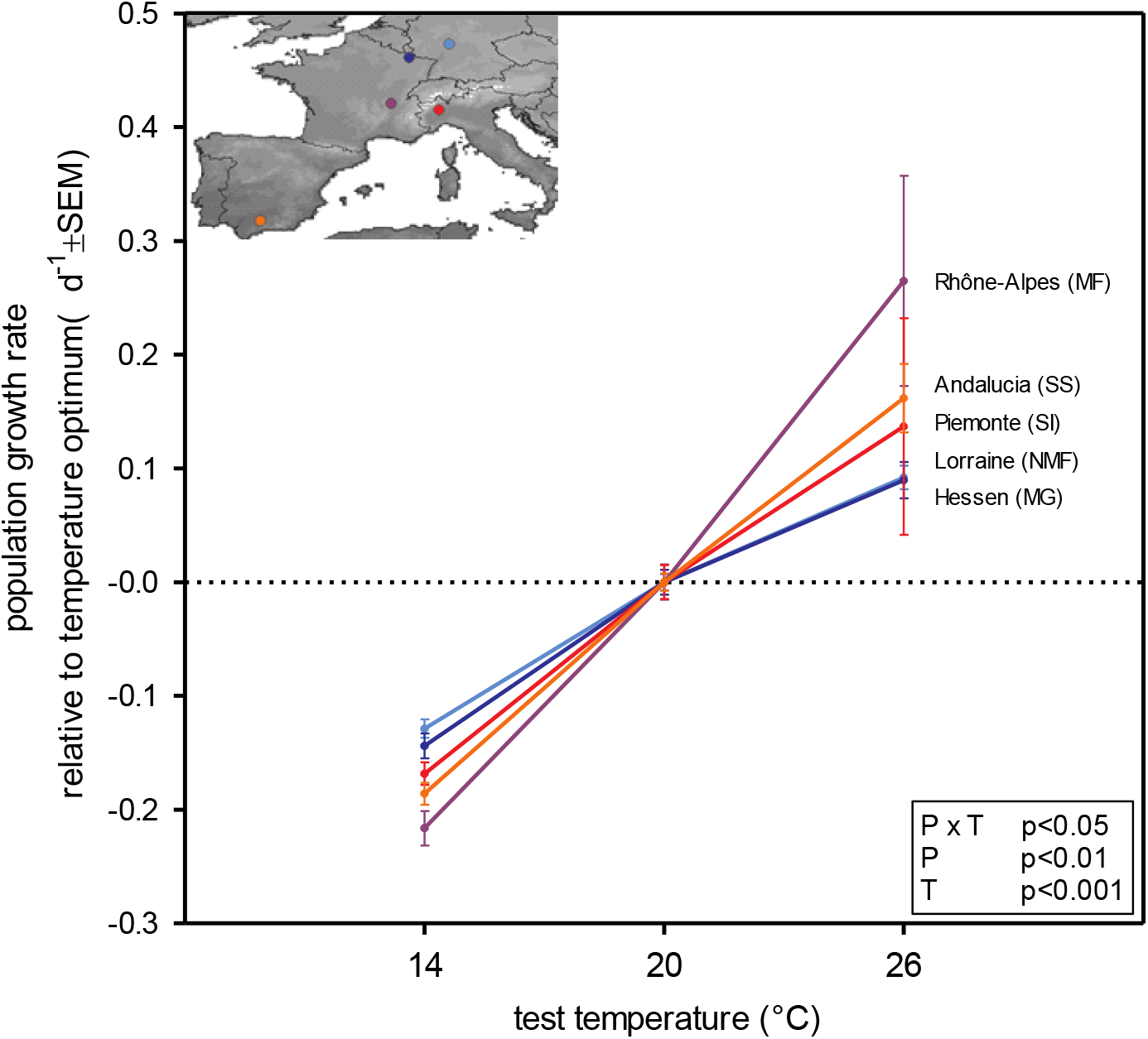
Population growth rate (PGR) of the natural *C. riparius* populations at different test temperatures. 20 °C constitute standard laboratory conditions and were chosen as optimal temperature to normalize the data for the tested populations, since we found overlapping variability (95 % CI) of the PGR in all populations. Locations of the populations along the European climate gradient are shown on the integrated map with the respective colour code. P-value thresholds of two-way ANOVA are given in the box: effect of population (P, F = 4.782), temperature (T, F = 105.4), and interaction of both factors (P × T, F = 2.418).

The differences in PGR between the five populations were based on different underlying life-cycle parameters for warm and cold test temperatures. At 14 °C mortality was significantly lower in the two Northern populations compared to the three remaining populations (Supporting Figure S3.2-A and Table S3.1-A). Furthermore, the two Northern populations showed a tendency of producing more fertile clutches per female at 14 °C than their relatives from more Southern locations (Supporting Figure S3.2-C and Table S3.1-C). These two parameters, however, showed increased variability at 26 °C and there was no clinal pattern. The number of eggs per clutch did not differ between temperatures or treatments (data not shown).

As expected for an ectothermic species, temperature had a significant effect on the PGR of the populations and accounted for 68 % of the total variance (F = 105.37, p < 0.0001). This effect was mainly driven by the temperature-dependent development of the larvae (Supporting Figure S3.2-B, Table S3.1-B). Furthermore, the PGR at the three test temperatures differed significantly between populations explaining 6.2 % (F = 4.78, p < 0.002) of the total variance, as well as the combined effect of temperature and population (6.3 % of the total variance, F = 2.42, p < 0.025).

### Demographic population history and migration

After trimming, the coalescence rate estimates from MSMC2 according to our quality threshold of a minimum of ten rate estimates per time slice (Supporting information 4), the informative time horizon for the inference of the demographic population history stretched from approximately 1,000 to 150,000 generations in the past. Based on a mean heterozygosity of 0.00426 and a switch error rate (SER) of 2.1 %, haplotypes of a mean length of 11,746 bases are informative for this analysis in *C. riparius*. This results in a tMRCA of 2,027 generations, giving a similar threshold of coalescence resolution in the recent past as derived after quality trimming coalescence rates.

In the late past, i.e. approximately 150,000 generations ago, N_e_ was estimated to an average of about 3.7 × 10^4^ individuals (Fig. 3A). During these generations the relative cross-coalescence between population-pairs revealed a good mixture of all populations with only a slight divergence drop about 100.0 generations ago (Fig. 3B). This ancestral population experienced a threefold decrease in Ne until 10,000 generations ago and by then the populations split during the following 9,000 generations. More or less complete population separation is estimated to have been established 1,000 generations ago. During time of population separation until 1,000 generations ago, N_e_ constantly increased reaching an average effective population size of 1.17 × 10^5^. Estimations for younger time points are ambiguous due to the limit of resolution of the analysis.

**Figure 3:**
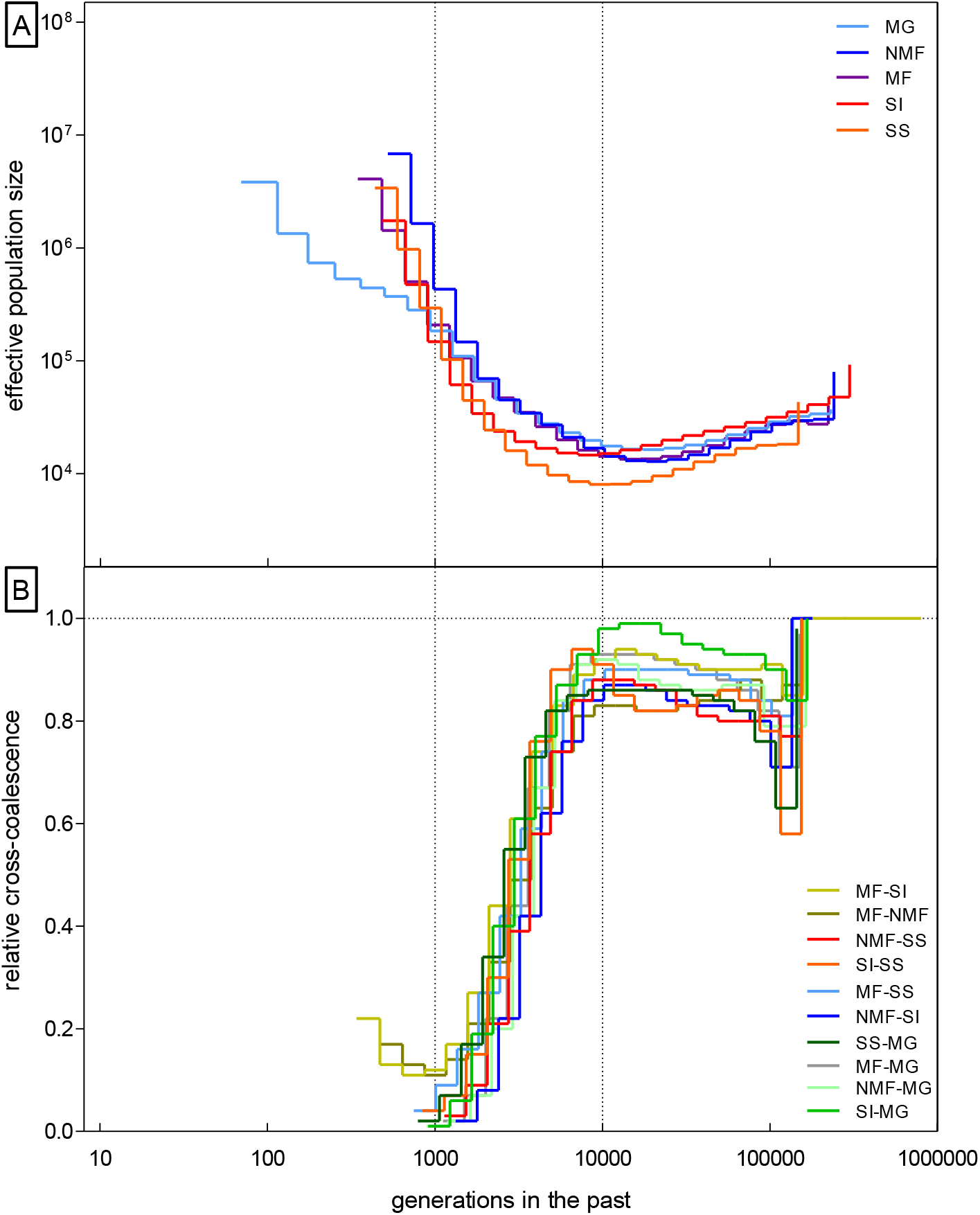
Results from genome-wide MSMC with eight individual haplotypes per population. (A) Effective population size and (B) relative cross-coalescence as indicator for population separation. To account for possible phasing error affecting the resolution, we only retained time indices with a minimum of ten coalescence rates.

The gene flow estimation integrated over the time period available for the analysis, revealed high migration rates, ranging from 3 × 10^−5^ to 9 × 10^−4^ (Supporting Table S6.1). Migration tended to radiate from the geographically centred population MF (Lyon), with less back migration. Least migration occurred across the Alps between Southern France and Italy.

### Neutral divergence threshold

Differentiation among population pairs calculated from 105,022 1 kb-windows along the genome was very low, with mean F_ST_ ranging from 0.032 to 0.111 (Supporting Table S6.2). Distribution of pairwise F_ST_ in these windows showed a weak isolation-by-distance pattern between populations (Fig. 4), though insignificant in Mantel’s test (ade4 package, R-Core 2015, 999 random permutations, p = 0.21). Comparing the empirical data with the distribution from the simulated drift expectation reveals a distinct mismatch of the data sets, especially for more diverged population-pairs (Fig. 4, Supporting information 5).

**Figure 4:**
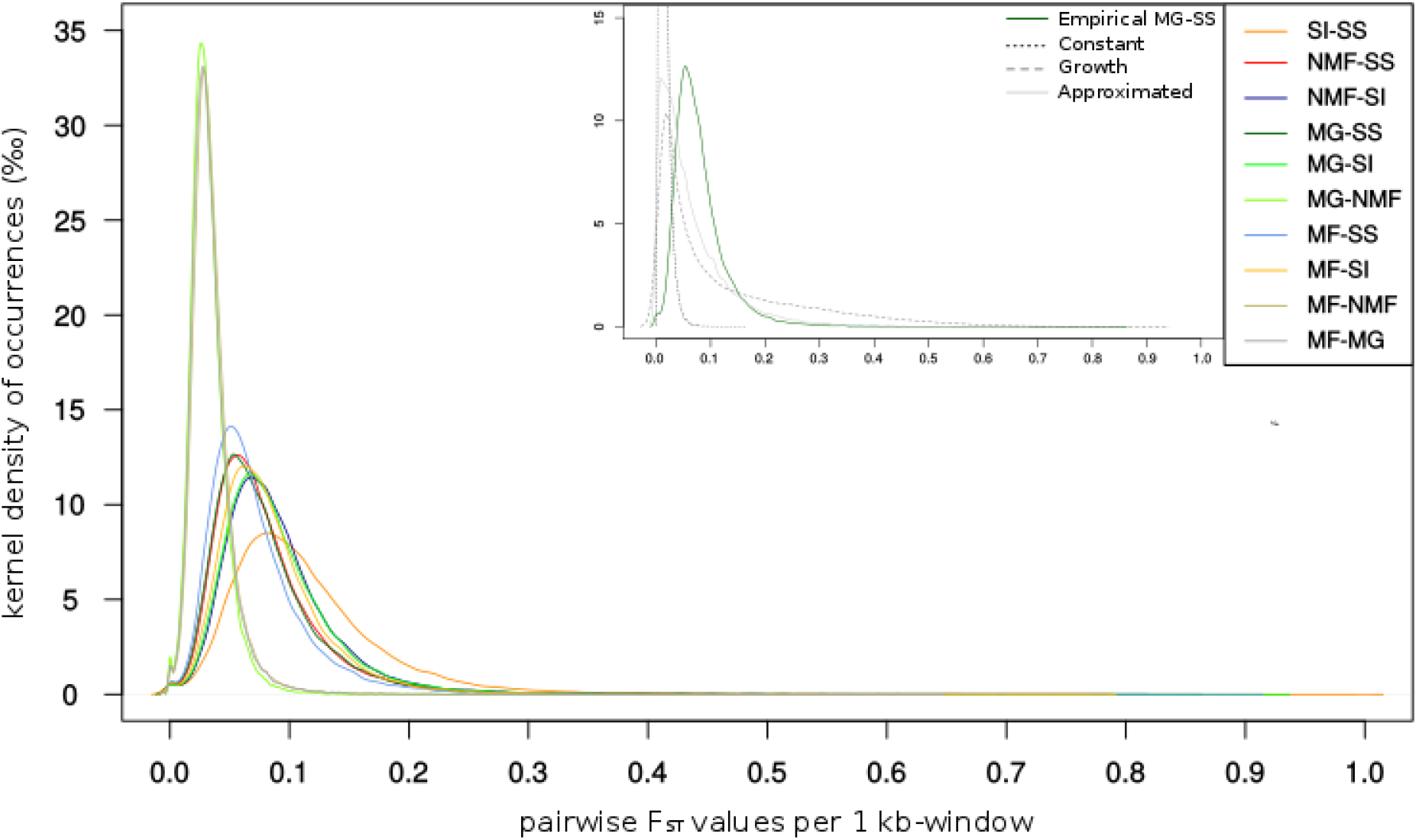
Distribution of pairwise F_ST_ per 1 kb-windows from *C. riparius* Pool-Seq data. Population comparisons are shown with different colours. The integrated graph shows one exemplary population-pair (MG-SS) in comparison to the respective simulated data from three different models.

The *constant* model clearly underestimated the differentiation among all populations, since the distributions peaked around zero. Observed empirical density functions were not comparably approximated by any model, i.e. even with increased complexity accounting for inferred population growth, adjusting for inferred population sizes and migration rates the *growth* and *approximated* model were unable to predict observed neutral divergence among populations.

We therefore decided for the overall more conservative upper 1 % tail of the empirical F_ST_ distribution as neutral divergence threshold for downstream analyses (see discussion below). Statistical F_ST_ thresholds of empirical data and those gained from the *approximated* model were most similar (Supporting Table S6.3). However, the statistical threshold from the empirical data was higher and thus more stringent in 7 out of 10 population comparisons. The retained statistical F_ST_ threshold (upper 1 % tail of the distribution) ranged from 0.1 to 0.36 for the respective population-pairs. A total of 4,360 highly diverged 1 kb-windows fell above the F_ST_ threshold from population comparisons (cf. Supporting Table S5.3 for pairwise numbers), of which 1,161 1 kb-windows were annotated to unique genes.

Joining adjacent 1 kb outlier windows of population-pairs to larger divergence regions resulted in 3,338 1 kb diverged windows above the 5 % F_ST_ cut-off, 1,711 of which could be joined to divergence regions larger than 2 kb. On average, a divergence region was 2478 bp long (+ 9764 bp) with a maximum length of 89 kb. Long divergence windows mostly occurred only once with the overall length distribution being strongly left skewed (Supporting Figure S6.1). The mean distance between divergence windows on the same scaffold was more than 80 kb (+ 151 kb, Supporting Figure S6.2).

### Partitioning among clinal and local adaptation

Environmental association analysis with LFMM was calculated on all 149,474 SNPs falling above the 99 % F_ST_ threshold of SNP-based genome-wide population comparisons (not to be confused with the 1 kb-window based population comparisons). Median z-scores were corrected with the estimated inflation factor λ for each of the three environmental variables (λp_recipitation_= 0.67, λ_temp_warm_= 0.71, λ_temp_cold_= 0.71). The three separate LFMM runs per climate variable resulted in 19,720 SNPs related to the precipitation gradient, 16,956 SNPs related to the gradient of warm temperatures, and 22,959 SNPs related to the gradient of cold temperatures (Fig. 5).

**Figure 5:**
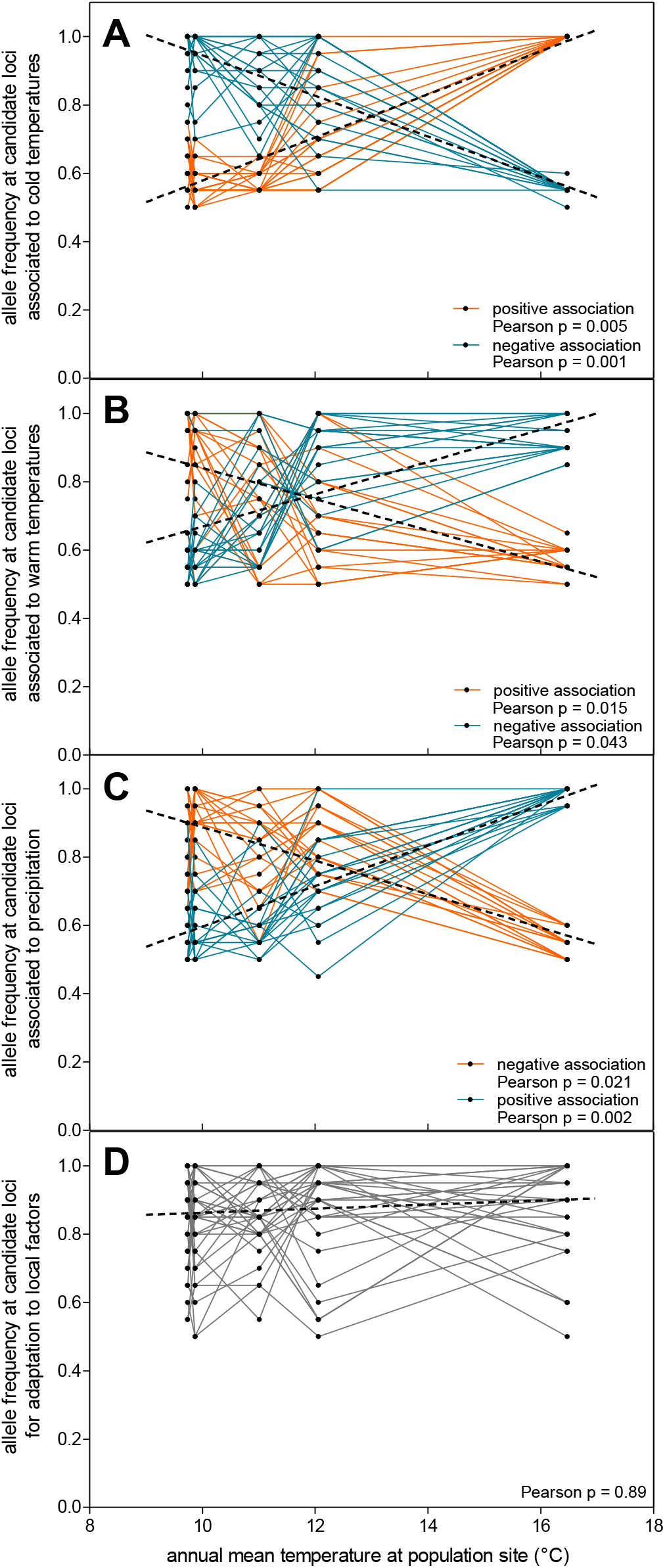
Correlation of *C. riparius* population growth rates from life-cycle experiments at three different test temperatures (14, 20, 26 °C) with annual mean temperatures at respective population sites as proxy for the climate gradient across Europe. Population growth rates at 26 °C show large variation, especially in populations that experience strong seasonal temperature differences (Rhône-Alpes, Piemont), thus, correlation shows a weak tendency. Population growth rates at 14 °C show a strong tendency in correlation to annual mean temperatures.

Annotation of clinal SNPs resulted in 1,551 unique genes as candidates for clinal adaptation: of those, 49 genes were private to the precipitation gradient, 196 genes private to the warm temperatures gradient, and 47 genes private to the cold temperatures gradient (Supporting Figure S8.1-A). The intersection of these candidates for clinal adaptation with those genes present within the annotated 1 kb-windows from the population comparisons (see above 1,161 genes) resulted in 162 genes (1.2 % of the annotated protein coding genes; Supporting Table S8.1; Appendix 1). These 162 clinal candidate genes have statistical support to be significantly diverged between at least two populations and were additionally found to be significantly correlated to one of the climate gradients (Supporting Figure S8.1-B). The intersection furthermore revealed 999 unique genes without clinal character (7.6 % of the annotated protein coding genes, Supporting Table S8.1; Appendix 1), as candidates for pure local adaptation.

Based on these different sets of candidate genes, we performed the GO term enrichment analysis to gain insight in biological processes that are likely to be involved in the adaptive evolution to local factors or climate factors (Fig. 6). Among the 999 genes putatively associated with local adaptation, the number of significantly enriched GO terms varied among populations from six in the Rhône-Alpes population (MF) to 19 in the Andalusian population (SS, Supporting Table S8.1). Associated to the 162 significant candidate genes for clinal adaptation, ten GO terms were found significantly enriched (Supporting Table S8.1), such as the terms ‘apoptotic process’ and ‘response to heat’. Among separated candidate genes, either associated to warm temperatures, cold temperatures, or precipitation, we found overrepresented GO terms that are private to the respective climate factor (Fig. 6, Supporting Table S8.1). KEGG pathways associated to clinal and local candidate genes were broadly overlapping (Appendix 1).

**Figure 6:**
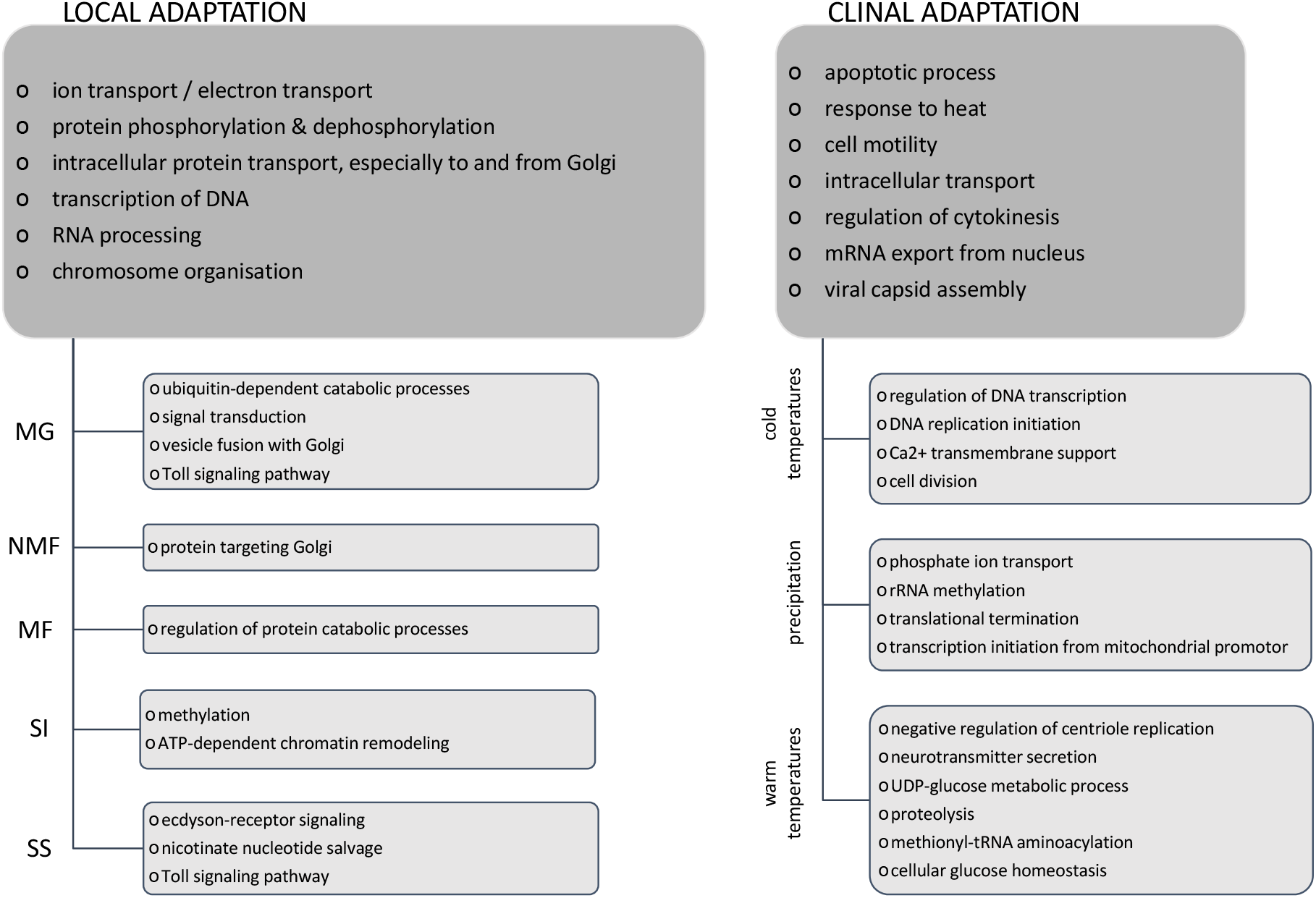
Functionally enriched GO terms associated to disentangled candidate genes for signatures of local adaptation in natural *C. riparius* populations (relevant and private GO terms shown, for complete lists cf. Supplementary Information S3) and for signatures of clinal adaptation along the climatic gradient across Europe. GO terms significantly enriched superior to the respective subgroups at top hierarchy, significantly enriched GO terms specific for subgroups below.

## Discussion

### Thermal adaptation of fitness-relevant traits in C. riparius

We experimentally demonstrated fitness-relevant thermal adaptation on the phenotypic level of five natural *C. riparius* populations sampled along a climatic gradient across Europe. According to the temperature conditions of their origin, population growth rates (PGR) of populations from colder sites in the North (annual mean temperature – AMT: 9.7-9.9°C) were higher at low test-temperatures compared to populations from warmer sites in the South (AMT: 12.1-16.5°C). The opposite was true for warm test-temperatures (Fig. 2, Supporting Fig. 3.1). This result is in line with previous findings of *C. riparius* populations from different locations among which the variability of PGR could be explained by a mixed effect of genetic drift, genetic diversity, and adaptation to the average temperature during the warmest month (Nemec *et al*. 2013). In *D. melanogaster*, heat and cold resistance differed between temperate and tropical populations (reviewed in Hoffmann & Weeks 2007) in the same clinal direction as found for *C. riparius* populations.

The contribution of traits underlying the PGR was different for the two extreme test-temperatures. Low temperatures significantly affected survival during the larval development and reduced viability of the offspring in warm-adapted populations, i.e. significantly less fertile clutches in at least two populations from warmer sites (Supporting Figure S3.2-C). High temperatures did not increase mortality in cold-adapted populations or directly affected their fertility, even though variability of these traits increased.

In *Drosophila*, heat knockdown resistance (time until flies are knocked down by heat in small tubes) and chill coma recovery (recovery time of flies after a cold shock) are the decisive phenotypic traits responding to heat and cold selection (e.g. Gilchrist *et al*. 1997; Hoffmann *et al*. 2002; Hoffmann & Watson 1993; Parson 1977). However, these traits can only indirectly be compared to population fitness. We here present a direct correlation of PGR, as integrated fitness measure, with the climatic origin of natural *C. riparius* populations and experimental test-temperatures. Such a correlation provides a meaningful model for the investigation of thermal adaptation on all levels (Hereford *et al*. 2004). Since our natural populations have been acclimatised to laboratory conditions for at most up to three generations, the differential response to thermal regimes in the experiments was most likely due to heritable differences among the natural populations, thus providing the necessary evidence to investigate the genomic footprint of this adaptation (Savolainen 2011).

### *Population genetic simulations underestimate neutral divergence in* C. riparius

Different evolutionary forces, i.e. selection, genetic drift, gene flow, and mutation, simultaneously shape the evolution of populations (Savolainen *et al*. 2013). Signatures of selection can therefore only be identified when patterns of neutral or endogenous evolution are subtracted from the signal of population differentiation, most importantly the impact of demography, migration, and potential endogenous selection barriers (Bierne *et al*. 2011; Flatt 2016). A previous study could show that a special transposable element might act as endogenous genetic barrier among the same European *C. riparius* populations as investigated here (Oppold *et al*. 2017). However, this pattern was not correlated to SNP differentiation between populations and can hence be neglected for this study.

A common and proven approach to infer selection among conspecific populations is to search for loci with increased population differentiation using F_ST_-based outlier tests (Narum & Hess 2011). Lewontin & Krakauer (1973) introduced the general idea that loci in which different alleles are selectively favoured in different populations should exhibit larger allele frequency differences between populations than purely neutrally evolving loci (Beaumont 2005). The challenge remains to identify a divergence threshold above which the influence of neutral processes that are not related to selection can reasonably be excluded. There is an ongoing debate that this threshold can either be derived in a model-based approach by simulating population divergence resulting from neutral processes (De Mita *et al*. 2013) or with a statistical threshold derived from the empirical F_ST_ distribution (Beaumont & Balding 2004). Population genetic modelling is used to infer divergence thresholds that account for species-specific population history. However, especially in non-model organisms the necessary population genetic parameters are scarce and their inference is challenging.

Using the genome-wide inference of N_e_ history (MSMC analysis, Fig. 3A), estimated migration rates between neighbouring populations (Supporting Table S6.1), and the spontaneous mutation rate of *C. riparius* (Oppold & Pfenninger 2017), we parameterised several species-specific population genetic models to simulate the actual drift-expectation in terms of F_ST_ distributions (Supporting information 5). Despite our substantial effort to obtain realistic empirical parameter estimates and therewith model neutral divergence among the populations, all applied models failed to reproduce the empirical F_ST_ distribution. The simulated data consistently underestimated population differentiation and their 99 % quantile F_ST_ thresholds were unrealistically low (Fig. 4, Supporting Table S6.3). The deviation between simulated and empirical data might be due to unresolved evolutionary circumstances in population history that could not be accounted for in our models. This could have several, mutually not exclusive reasons: (1) inadequacy of current coalescence/population genetic models for estimation of gene flow and Ne to account for differential evolutionary speed of populations; (2) more complex population history; (3) pervasive action of selection throughout the entire genome; (4) chromosomal inversion polymorphisms that disproportionately increase population differentiation; (5) cryptic introgression in parts of the species range. A more detailed discussion of these five points in the following:

1. The number of generations per year of a *C. riparius* population strongly depends on the temperature conditions at the respective location throughout the year (Oppold *et al*. 2016). This leads to substantially different ‘evolutionary speeds’ among populations of different temperature regions with up to 3-fold different numbers of generations per year. Population genetic consequences of this temperature dependence are, however, not accounted for in current population genetic models used e.g. for coalescence analysis of demography and migration. The coalescence approach of MSMC2 uses historical recombination events to infer past population sizes from the distribution of coalescence times (Ellegren & Galtier 2016). The inference is based on a uniform time index for coalescence events, i.e. an equal number instead of differential numbers of generations among populations across temperature gradients. It is currently not possible to correct for this and thus, coalescence of multivoltine insect populations along climate gradients is fundamentally biased. Results of the MSMC2 analysis for the inference of Ne include this bias as soon as populations started separating from each other, i.e. when they are expected to have experienced different temperature regimes (see discussion below). Ne estimates of the recent past and the timing of separation events might thus be incorrect. Similarly, results of the gene flow analysis are based on the distribution of SNPs among populations. Populations from warmer regions can accumulate more polymorphisms while passing more generations (Oppold *et al*. 2016), which in turn might lead to an inflated migration rate estimate towards populations of cooler temperature regions (see discussion below).
2. Complex and hierarchical population structures are known to inflate F_ST_ distributions with an excess of false significant loci (Excoffier *et al*. 2009), which could in principal explain the result of our population genetic modelling. The genome-wide inference of population history (MSMC2 analysis), however, indicated one ancestral panmictic *C. riparius* population that started diverging approximately 10,000 generations ago until populations reached almost complete separation 1,000 generations ago (Fig. 2B). As long as the coalescence analysis indicates a single panmictic population and early population separation, the above discussed influence of evolutionary speed on inference of demographic history should be neglectable. Upon this finding, we can reasonably assume that all populations derived from a single ancestral population without hierarchical substructuring. Nevertheless, due to the temperature dependence of generation time it is difficult to convert the scaling of generations to an absolute time scale in years, neither knowing the precise population location nor thermal conditions at that time. Based on previous modelling of the potential number of generations per year at thermal conditions during the last glacial maximum (Oppold *et al*. 2016), we can roughly assume five to eight generations per year for the ancestral panmictic population. This converts the period of population separation to a time horizon from 2000-1250 years ago until the last 200-125 years. Associated with the process of population separation, Ne has been increasing drastically. *C. riparius* is known for its broad ecological tolerance, yet larvae preferably inhabit rivers with organic pollution (Armitage *et al*. 1997; Groenendijk *et al*. 1998). The increase in N_e_ might therefore be correlated to the increase in human settlements over the last centuries (Antrop 2004) and anthropogenic impact (intensification of agriculture, industrialisation, increase in waste water) on the environment in general, and water bodies in particular. Migration analysis indicated that gene flow radiated from the current Lyon population towards the other populations, even though this result has to be regarded carefully due the above discussed fundamental bias of the estimation. Disregarding absolute migration rates, the general migration pattern suggests that an expansion across Europe might have originated from central France (Supporting Table S5.1). Since we can exclude a significant difference in N_e_ between populations (Fig. 3A), Tajima’s D analysis suggests that positive selection has been playing a major role in populations at the outer margins of the investigated climatic gradient (Supporting information Box 7, Figure S7.2). If populations expanded from central France, i.e. the centre of the cline, this might provide an explanation for this spatial pattern of positive selection. According to the allele surfing phenomenon, mutations occurring at the edge of the range expansion are lost at a reduced rate and can more easily be driven to fixation (Klopfstein *et al*. 2006). Therefore, the time of population range expansion is an evolutionary important period, where mutations can accumulate and contribute to adaptation processes. By the time population separation was completed, we can assume that our *C. riparius* populations had established their current geographic distribution across Europe, inevitably experiencing different temperature regimes. Demography estimates from the recent past therefore include the above described bias and cannot be resolved due to differential evolutionary speed among populations. In general, this adds a cautionary note to the analysis of historical population history in multivoltine ectotherms with a wide distribution range.
3. Pervasiveness of selection could, in principle, explain the observed discrepancy between simulated and empirical data. In that case, local selection of polygenic traits would result in a shift of F_ST_ distributions to higher values. Evidence for such a scenario comes from investigations comparing *Drosophila* species suggesting that most of the genome is under selection (Sella *et al*. 2009). However, our study compared intraspecific *C. riparius* populations with high gene flow (assumed from high migration rates and low population differentiation with a mean F_ST_=0.07 among all populations) and therefore pervasive selective divergence appears rather unlikely.
4. There are many cases of inversion polymorphisms with clinal frequency patterns described in *Drosophila* that are correlated to environmental factors or even seasonal climate variations (Kapun *et al*. 2016). Such structural mutations can lead to strong signals of divergence along the inverted sequence between populations. Inversions were described also for many *Chironomus* species, although such polymorphisms were hardly observed in *C. riparius* (then called *C. thummi*, Acton 1956). Furthermore, divergent regions among *C. riparius* are on average rather short (2,478 bp; Supplementary Figure S5.1) and widely distributed (on average 80 kb; Supplementary Figure S5.2), whereas inversions are expected to span large regions along chromosomes (Caceres *et al*. 1997). It is therefore unlikely, though cannot be entirely ruled out, that inversions are involved in the clinal differentiation or even enhance thermal adaptation in *C. riparius* similarly as documented for *Drosophila*.
5. A scenario of differential introgression from related *Chironomus* species appears possible. Whole genome sequencing revealed reticulated evolution across the entire species tree, and hybridization with introgression among related multicellular eukaryotic species seems to be pervasive (Mallet *et al*. 2016; Shapiro *et al*. 2016). Hybridization between *Heliconius* species appears to be a natural phenomenon and Mallet *et al*. (2007) even deduced that introgression may often contribute to adaptive evolution. The genus *Chironomus* is very species-rich and more than 4,000 species are unambiguously aquatic during larval development, often dominating aquatic insect communities (Ferrington 2008), such that larvae of many different species can co-occur at the same location (Pfenninger et al. 2007). The closest relative to *C. riparius* is the morphologically cryptic sister-species *C. piger* that shares the same habitats (Schmidt *et al*. 2013). *C. piger* is described to occur in Northern and Central Europe (Grebenjuk & Tomilina 2014; Ilkova *et al*. 2007; Michailova *et al*. 2015; Pedrosa *et al*. 2017b; Sokolova *et al*. 1992), widely overlapping with the *C. riparius* distribution range. During sampling for this study, we also found co-occurrence of the two species at the Northern-most sampling sites (MG and NMF) and the discrepancy between empirical and simulated F_ST_ distributions was especially large in comparisons with those two populations. Hybridisation of the sister-species is possible under laboratory conditions (Keyl 1963 and own observations), though previous studies found little evidence for ongoing hybridisation in the field (Hägele 1999; Pedrosa *et al*. 2017b; Pfenninger & Nowak 2008; Pfenninger *et al*. 2007). Experimental analysis of niche segregation between the sister-species showed that *C. riparius* had a higher fitness at higher constant temperatures and larger daily temperature ranges (Nemec *et al*. 2012). A potential differential *C. piger* introgression along the cline of *C. riparius* distribution can thus not be excluded, and requires deeper investigation.

Since it was, for the above potential reasons, not possible to derive a reasonable model-based neutral divergence threshold, we resorted to a statistical threshold as advocated e.g. by Beaumont (2005). This threshold was in addition more conservative (i.e. higher F_ST_) for the majority of pairwise comparisons (Supporting Table S6.3).

### Functional basis of local and clinal adaptation in C. riparius

A population genetic bottom-up approach including pooled sequencing and a high quality genome annotation is powerful to build new hypotheses about genotype-phenotype interactions supported by genomic signatures of positive selection (Kolaczkowski *et al*. 2011). By overlapping gene lists of the F_ST_ outlier approach with gene lists of the environmental association analysis (EAA), it was possible to disentangle signatures of local and clinal adaptation. EAA revealed candidates of clinal climate adaptation, i.e. SNPs that showed gradual allele frequency changes in correlation to the gradual change of one specific climate variable (Fig. 5). The rationale behind this is that climatic gradients constitute predictable continuous clines that enable the investigation of clinal adaptation if genotypic variation correlates with the gradual pattern (Adrion *et al*. 2015). While climate factors produce a spatially continuous selection regime, there are likely additional local selection pressures that act on populations only in their specific habitat. Thus, there are SNPs without association to climate variables (Fig. 5) that are still highly divergent at least among some populations (defined as the upper 1 % tail of the site-specific F_ST_ distribution). These SNPs are therefore considered as candidates of local, non-clinal adaptation.

To identify possible selective forces driving local adaptation (i.e. non-clinal), one needs to consider the ecological niche. *C. riparius* larvae inhabit small ditches and streams with increased carriage of organic matter, mostly in agricultural areas or streams receiving waste effluents from urban areas (Armitage *et al*. 1997; Calle-Martínez & Casas 2006). Consequently, *C. riparius* populations need to adapt to their local pollutants, i.e. pesticides (Müller *et al*. 2012), metals (Pedrosa *et al*. 2017a; Wai *et al*. 2013), organic pollution (Vogt *et al*. 2007), and its effects on physicochemical conditions of the water body as decreased oxygen content or hydrogen sulphide levels. Results of the functional enrichment analysis indicate that candidates of local adaptation are significantly enriched for genes that can be associated to detoxification processes, as for example transport processes, phosphorylation, epigenetic response, immunity, and larval development (Tab. 1, Fig. 6). Some of these GO terms were also previously identified in studies with *D. melanogaster* (Tab. 1; Fabian *et al*. 2012; Kolaczkowski *et al*. 2011; Levine & Begun 2008). Those studies actually aimed at investigating patterns of clinal differentiation among populations, whereas we found the same GO terms to be significantly enriched among candidates of non-clinal/local adaptation. Reasons for this discrepancy might be that either only the endpoints of a cline were investigated (Levine & Begun 2008; Turner *et al*. 2008), or clinality of SNPs was examined by manually classifying allele frequency changes between three populations across latitudes (Fabian *et al*. 2012). The comparison of only two *Drosophila* populations, however, lacks resolution to disentangle signatures resulting from site-specific local or clinally varying selection pressures. Moreover, the advantage of using EAA that are based on Bayesian methods, as the here applied LFMM, is that confounding effects can be corrected for (Frichot & Francois 2015) which is not possible when allele frequency changes of separate SNPs are categorized. Bergland *et al*. (2016) suggested that instead of spatially varying selection pressures, demographic admixture events from two ancestral *Drosophila* populations generated the clinal genetic variation at approximately one third of all common SNPs across clines on different continents. This highlights the importance of accounting for confounding genetic effects due to population structure.

**Table 1:**
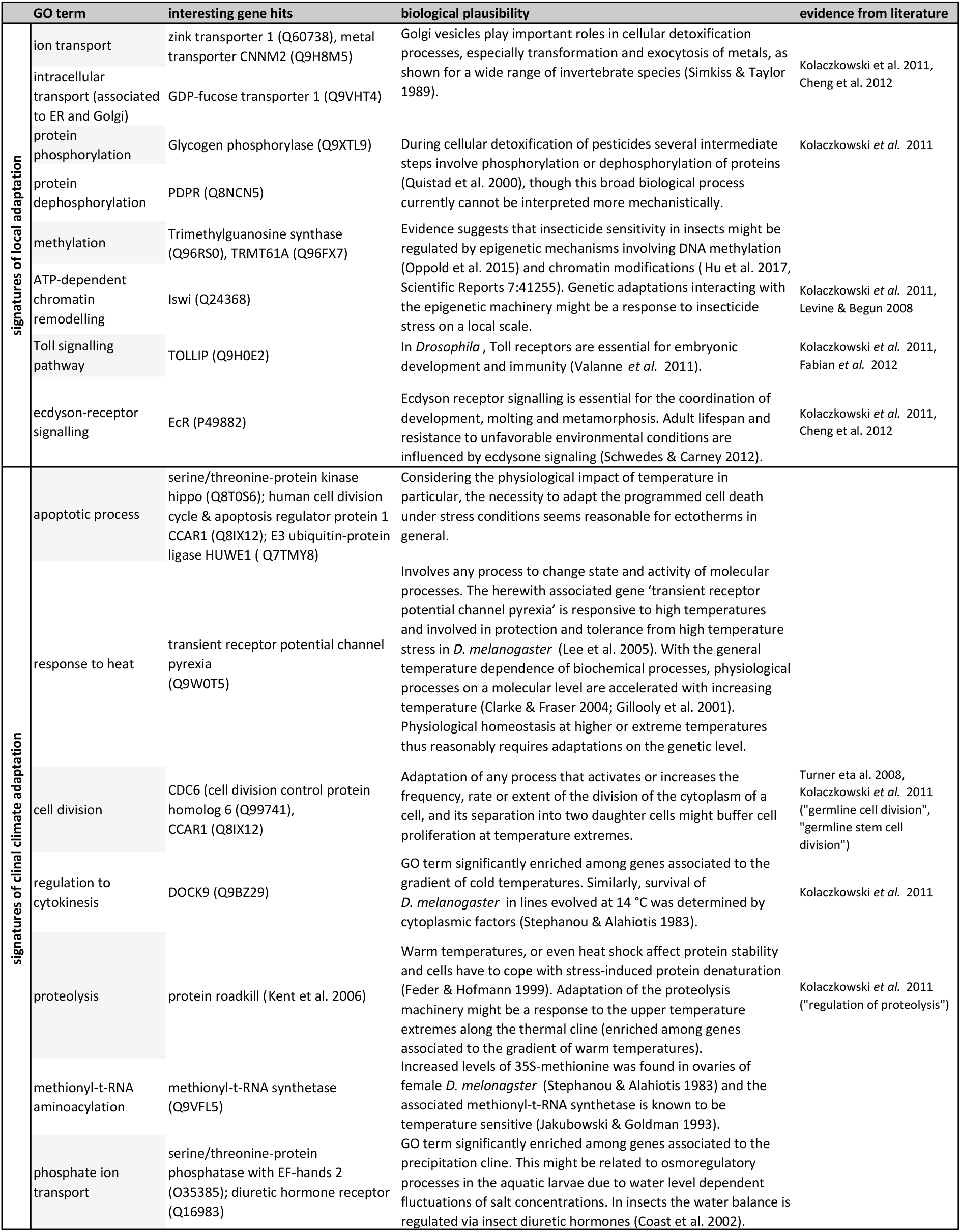
Detailed discussion of relevant GO terms that were significantly enriched among candidate genes of local or climate adaptation. Underlying gene hits are given with UniProt accession numbers. Literature reference is given if the same or closely related GO term was previously identified in insects.

Nevertheless, the here significantly enriched GO terms among candidates of local adaptation were also identified to be relevant for adaptive processes in other insects, suggesting that these biological functions seem to be ecologically relevant in *C. riparius*, *Drosophila* (Fabian *et al*. 2012; Kolaczkowski *et al*. 2011; Levine & Begun 2008), as well as *Anopheles gambiae* (Cheng *et al*. 2012).

The intersection of candidate genes resulting from F_ST_ outlier analysis and the EAA resulted in 162 candidate genes for climate adaptation that were significantly diverged among populations as well as significantly correlated to one climatic cline (Supporting Fig. S8.1). Therefore, 1.2 % of all protein coding genes show signatures of climatic selection (Supporting Table S8.1). Among these, we found significant enrichment for genes associated with apoptosis, response to heat, cell division, regulation of cytokinesis, and proteolysis (Tab. 1, Fig. 6). The three latter GO terms (or closely related ones) were also found significantly enriched among candidate genes of clinal differentiation in *Drosophila* (Kolaczkowski *et al*. 2011; Turner *et al*. 2008). They all describe fundamental biological processes that are important for cell survival, suggesting that especially climate extremes majorly drive climate adaptation.

To increase resolution on the potential functional basis of climate adaptation, we considered the three climate variables separately. This demonstrated that different biological processes seem to be involved in adaptation to the gradient of cold temperatures, warm temperatures or precipitation (Fig. 6). Further investigations on the level of molecular functions will be necessary for a fine scale comparison to previously described functional networks, as e.g. the candidate genes network of cold tolerance in *Drosophila* (Bozicevic *et al*. 2016). Among precipitation candidate genes we found significant enrichment of phosphate ion transport with the diuretic hormone receptor as key gene (Tab. 1). Competition strength of *C. riparius* larvae against the co-occurring sister species *C. piger* was found to be correlated to precipitation of the warmest quarter/month and in particular negatively correlated to water conductivity, nitrate, and calcium carbonate (Pfenninger & Nowak 2008). This indicates the relevant fitness effect of water composition, as direct result of varying precipitation levels, for *C. riparius* larvae and thus driving adaptation along the precipitation cline.

Functional enrichment analyses and the categorisation of gene ontology come along with the problem that the interpretation of results finally relies on biological interpretation of plausibility and relation to literature (Pavlidis *et al*. 2012). Especially, functional clustering of genes along chromosomes and/or evolutionary population histories may eventually cause over-representation of biological categories, similar as driven by positive selection (Pavlidis *et al*. 2012). In our study however, the influence of functional clustering of candidate genes should be neglectable since highly differentiated regions of exceptional population differentiation were found to be rather short and widely distributed across the genome (Supporting Figure S6.1-2). Our divergence regions with on average 2,478 bp are not expected to span over multiple genes with on average 4.7 exons at an average length of 355 bp in *C. riparius* (Oppold *et al*. 2017). Moreover, population history should not produce false positive GO enrichment, since the demographic analysis with our *C. riparius* populations did not reveal drastic bottlenecks that could have produced enrichment patterns similar to positive selection. Considering the relevance of polygenic traits (Wellenreuther & Hansson 2016), the number of false negative divergence regions might be high since we have chosen a rather conservative F_ST_ threshold. The here presented results can therefore be considered as robust and most prominent signatures of adaptation among *C. riparius* populations, although revealing only a fraction of the complete genomic footprint. We found several significantly enriched GO terms with biological plausibility indicative for either local adaptation or clinal climate adaptation (Tab. 1 and Appendix 1), providing a promising basis for their validation in downstream analyses. Candidate genes for local as well as clinal adaptation showed a large overlap in the associated KEGG pathways (Supporting Table S781, Appendix 1). Those involved metabolism of carbon, nitrogen, methane, and sulphur pathways. Even though there are clear differences on the gene level signatures of local and clinal adaptation do not show a clear separation on the complex level of molecular pathways.

### Conclusions

Climate poses a fundamental influence on evolutionary dynamics of multivoltine ectotherms, which becomes apparent in population genetic modelling of their neutral genetic divergence. With our integrative analysis on the phenotypic and genotypic level, we were able to separate the footprints of clinal climate adaptation from habitat-specific local adaptation as well as from neutral evolution among *C. riparius* populations. The comparison of the investigated natural populations revealed the impact of adaptive evolution despite high levels of gene flow and thus on average weak population differentiation. Important biological processes overlapping with findings in other studies with different insect species were identified to be involved in climate adaptation. Deeper investigations on the gene level will be necessary to clarify the decisive genetic factors that underlie those adapted biological processes.

## Acknowledgement

We thank Stephan Schiffels and Olivier Francois for help and recommendations on the MSMC2 analysis and LFMM analysis, respectively. We also thank Bob O’Hara for comments and support with the population genetic modelling. Four anonymous reviewers provided constructive recommendations for improving the manuscript. Funding was provided by DFG (PF390/8-1). Ann-Marie Waldvogel acknowledges funding by a scholarship of the FAZIT-Stiftung.

## Data accessibility

Raw data from whole genome individual resequencing available at European Nucleotide Archive (ENA project number pending).

## Authors’ contribution

M.P., T.H., and A.-M.W. conceived the study; A.-M.W. performed common-garden experiments, prepared sequencing of individual resequencing data, phased individual resequencing data and performed MSMC2, and analysed Pool-Seq data; A.-M.W. and M.P. performed the Tajima’s D analysis, A.W. and M.P. performed population genetic modelling; A.-M.W. and A.W. performed environmental association analysis; M.P. performed gene flow analysis; B.F. performed functional enrichment analysis; T.S. provided bioinformatic support and custom scripts; S.P. supported Pool-Seq analyses; H.S. supported genomic analyses; A.-M.W., A.W., M.P., B.F., H.S., S.P., and T.H. drafted the manuscript.

